# Testing structural identifiability by a simple scaling method

**DOI:** 10.1101/2020.02.04.933630

**Authors:** Mario Castro, Rob J. de Boer

## Abstract

Successful mathematical modeling of biological processes relies on the expertise of the modeler to capture the essential mechanisms in the process at hand and on the ability to extract useful information from empirical data. The very structure of the model limits the ability to infer numerical values for the parameters, a concept referred to as structural identifiability. Most of the available methods to test the structural identifiability of a model are either too complex mathematically for the general practitioner to be applied, or require involved calculations or numerical computation for complex non-linear models. In this work, we present a new analytical method to test structural identifiability of models based on ordinary differential equations, based on the invariance of the equations under the scaling transformation of its parameters. The method is based on rigorous mathematical results but it is easy and quick to apply, even to test the identifiability of sophisticated highly non-linear models. We illustrate our method by example and compare its performance with other existing methods in the literature.

**Author summary:** Theoretical Biology is a useful approach to explain, generate hypotheses, or discriminate among competing theories. A well-formulated model has to be complex enough to capture the relevant mechanisms of the problem, and simple enough to be fitted to data. Structural identifiability tests aim to recognize, in advance, if the structure of the model allows parameter fitting even with unlimited high-quality data. Available methods require advanced mathematical skills, or are too costly for high-dimensional non-linear models. We propose an analytical method based on scale invariance of the equations. It provides definite answers to the structural identifiability problem while being simple enough to be performed in a few lines of calculations without any computational aid. It favorably compares with other existing methods.

## Introduction

Mathematical models contribute to our understanding of Biology in several ways ranging from the quantification of biological processes to reconciling conflicting experiments [1]. In many cases, this requires formulating a mathematical model and extracting quantitative estimates of its parameters from the experimental data. Parameters are typically unknown constants that change the behavior of the model. While it is usually recognized that parameter estimation requires the availability of sufficient informative data, sometimes it is not possible to estimate all parameters due to the structure of the model (whatever the quantity or quality of the data). This inability is referred to as ‘structural identifiability’, a concept introduced decades ago by Bellman and Åström [2, 3], as opposed to the ‘practical identifiability’ that depends on limitations set by the data. Practical identifiability has important consequences that can lead to questionable interpretations of the data leading to some recent controversy around this point [4, 5]. Structural identifiability is a necessary condition for model fitting and should be used before any attempt to extract information about the parameters, and as a test of the applicability of the model itself. Structural identifiability can be qualified as global or local. Global structural identifiability tests the ability to estimate unique sets of parameters, while local (or simply, structural identifiability) means that parameters can be estimated only in a limited subset of the space of parameters, *i.e.*, only combinations of parameters are identifiable [6–10]. In practical terms, these definitions can be translated into the language of sensitivity analysis as identifiability requires that (i) the columns of the sensitivity (or, equivalently, the elasticity) matrix are linearly independent, and (ii) each of its columns has at least one large entry [11, 12]. Traditionally, work primarily focused on linear systems [2, 3, 13] based on ordinary differential equations (ODE). For non-linear models, those methods cannot be applied, so many methods have been proposed in the literature to address structural identifiability. Early attempts were based on power series expansions of the original non-linear system [14], the similarity transformation method [15–17] or the so-called direct-test method proposed by Denis-Vidal and Joly-Blanchard [18, 19]. These methods exploit the definition of identifiability either analytically [18] or numerically [20–25], but they are not generically suitable for high-dimensional problems. Xia and Moog [6, 26] proposed an alternative to these classical methods based on the implicit function theorem, but this method also becomes involved to apply for complex models [27]. Another approach that is becoming mainstream is based on the framework of differential algebra [28–31]. These methods are also difficult to apply, requiring advanced mathematical skills and, in some cases, replace highly non-linear terms by polynomial approximations that simplify the analysis. On the positive side, they are based on rigorous mathematical theories, are suitable for non-linear models and, more importantly, they can be coded using existing symbolic computational libraries. In this regard, it is worth mentioning DAISY [32], GenSSI [33], COMBOS [34] or, more recently, SIAN [35].

In almost all cases, the major disadvantage of these methods is their difficulty to apply them to even a few differential equations, hence requiring advanced mathematical skills and/or dedicated numerical or symbolic software (that is frequently unable to handle the complexity of the problem). This explains why, despite the huge volume of publications in the field of theoretical biology, only a few address parameter identifiability explicitly. In this paper, we introduce a simple method to assess local structural identifiability of ODE models that reduces the complexity of existing methods and can bring identifiability testing to a broader audience. Our method is based on simple scaling transformations, and the solution of simple sparse systems of equations. Identifiability for stochastic models [36] is out of the scope of our work.

## Method

### A couple of motivating examples

Consider a simple *death* model in which the death rate is the product of two parameters λ_1_ and λ_2_, namely

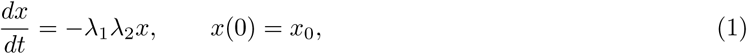

with the solution

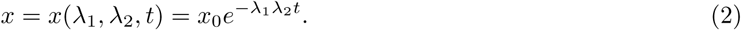

It is evident that from an experiment only the product λ_1_λ_2_ can be inferred, and not any of the two independently. Following the ‘actionable’ definition in Ref. [11], local structural identifiability is directly linked to the linear independence of the columns of the elasticity (or, sensitivity) matrix. In this case, the elasticity matrix would be simply a 1 × 2 matrix,

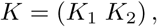

with

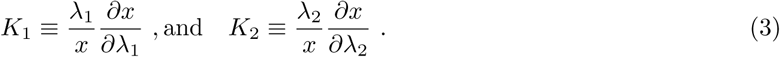

We now propose to multiply λ_1_ with a generic scale factor *u*, and to divide λ_2_ by the same factor, such that the solution remains invariant. Deriving the scaled solution of eq. (2) with respect to that scale factor *u*, and by the chain rule,

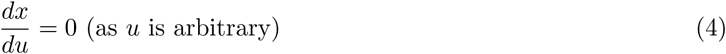

and, also,

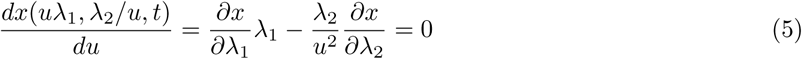

where the last equality follows from Eq. (4)

Rearranging Eq. (5) and dividing by *x*,

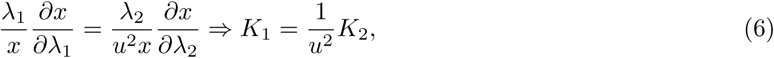

so both *columns* of the elasticity matrix are linearly dependent and, accordingly, λ_1_ and λ_2_ are unidentifiable. In this case we had complete knowledge of the solution, and consequently, it was straightforward to find the right way to introduce the scaling *u*. Fortunately, this simple scaling calculation can also be performed directly on eq. (1). Introducing two unknown scaling factors, *u*_1_ and *u*_2_, into that equation,

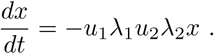

Requiring that this remains identical (or, more formally, *invariant*) to Eq. (1), i.e., λ_1_λ_2_*x* = *u*_1_λ_1_*u*_2_λ_2_*x*, hence *u*_1_*u*_2_ = 1. The fact that *u*_1_ and *u*_2_ cannot be solved individually, also means that the real values of λ_1_ and λ_2_ cannot be determined, namely both parameters are unidentifiable.

Next consider a death model with immigration:

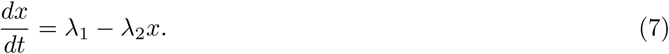

In this case, to leave the system invariant we need to find *u*_1_ and *u*_2_ such that

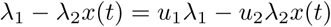

for all values of *x* at any time. Rearranging the latter equation,

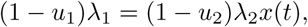

where the left-hand side of the last equation is a constant and the right-hand side depends on time. Hence the only possible solution to the latter equation is *u*_1_ = *u*_2_ = 1 implying that both λ_1_ and λ_2_ are locally identifiabile. Notice the difference with the preceding case, Eq. (1), in which an infinite number of combinations of the scaling factors satisfy the invariance condition.

These simple examples illustrate how scaling invariance of the model equations can be used to determine whether the parameters are unidentifiable or not. We prove this result more rigorously in the Supporting Information.

### Description of the method

Let us define a general ODE model characterized by the time evolution of *n* variables, *x*_*i*_(*t*), depending on *m* parameters λ_*j*_,

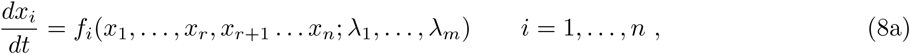

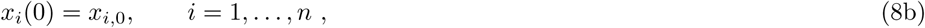

where the functions *f*_*i*_ depend on the specific details of the problem at hand and *x*_*i*,0_ are the initial conditions. We need to distinguish between those variables that can be observed (measured) in the experiment, *x*_1_… *x*_*r*_, and those which cannot (they are often referred to as *latent* variables), *x*_*r*+1_… *x*_*n*_.

According to Sec. 1 in the Supporting Information, each function *f*_*i*_ is split into *M* functional independent *summands*, *f*_*ik*_,

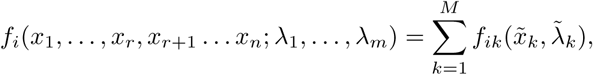

having the property that *f*_*ik*_ is functionally independent of *f*_*il*_ for every *k* ≠ *l*. For brevity, 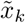, 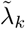 denote the subset of variables and parameters included in the function *f*_*ik*_. Typical examples of functionally independent functions are *f*_11_ = *ax*_1_, *f*_12_ = *bx*_1_*x*_3_, *f*_13_ = (*c* + *x*_4_)^−1^, and examples of dependent functions would be

##### Box 1: Summary of the scale invariance local structural identifiability method introduced in this work.

1. Scale all parameters and all unobserved variables by unknown scaling factors, *u*:

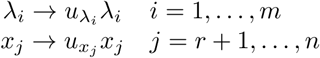

and substitute them into Eqs. (10) below.
2. Equate each functionally independent function, *f*_*ik*_, to its scaled version. Namely,

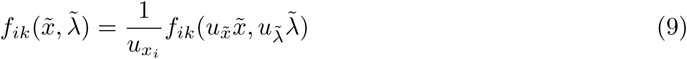

where 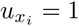 for 1 ≤ *i* ≤ *r* and the prefactor in the right-hand side of the equation comes from the scaling of 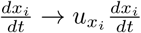. From Eq. (8b) it follows that 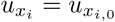.
3. From the Eqs. (9), find combinations of the scaling factors *u* that leave the system invariant. Hereafter, we will denote these as the **identifiability equations** of the model.
4. Only the parameters λ_*i*_ with a solution *u*_*i*_ = 1 are identifiable.

*f*_11_ = *ax*_1_*x*_2_ and *f*_12_ = *bx*_1_*x*_2_ (see Supporting Information for details). Note that it is not required that *f*_*ij*_ and *f*_*kj*_ are independent (as they appear in different equations). For instance, in the example in Eq. (7)

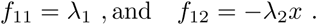

We summarize our method in Box 1.

In summary, our method reduces the complexity of finding identifiable parameters to finding which scaling factors do not satisfy the trivial solution *u*_*i*_ = 1. In the literature, when a scaling factor is related to one of the latent variables *x*_*r*+1_… *x*_*n*_, if 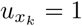, then *x*_*k*_ is said to be *observable* [10]. Thus, our method addresses at the same time identifiability and observability. Additionally, irreducible equations involving two or more parameters provide the so-called identifiable groups of variables that cannot be fitted independently. In the case of the pure death model above, the identifiability equation 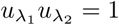 is a signature of the unidentifiable group λ_1_λ_2_. This is interesting as groups involving latent variables (for instance, 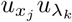) would inform future experiments aimed to observe that variable and decouple that group.

It is also worth mentioning that our identifiability test (illustrated by example in the Supporting Information) provides a simple way to find a type of symmetry that is related to scale invariance. More sophisticated methods have been introduced in the literature to address other symmetries [37–39] using the theory of Lie group transformations, however, that approach involves complex calculations assisted by symbolic computations.

## Results

### The main result

Consider a model described by a set of *n* ordinary differential equations (ODE)

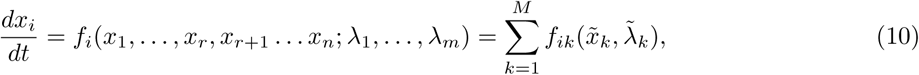

where *f*_*ik*_ is functionally independent of *f*_*il*_ for every *k* ≠ *l* (namely, they satisfy the generalized Wronskian theorem; see the Supporting Information). For the sake of simplicity, we denote 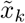 and 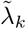 the subset of variables and parameters of function *f*_*ik*_.

Motivated by Eqs. (1)–(3), we seek for scaling of the parameters that leave the system invariant. As we prove below, this invariance (or lack of) is related to the identifiability of the parameters. Hence, if we define the following scaling transformation:

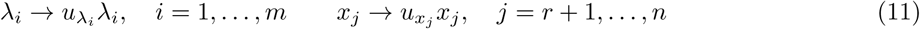

(where the variables *x*_1_… *x*_*r*_ are unmodified as we can measure them in the experiment) we can write the following set of re-scaled equations:

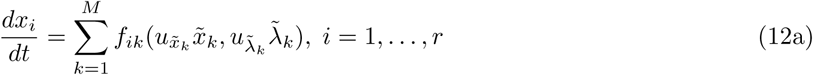

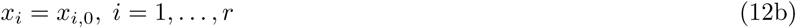

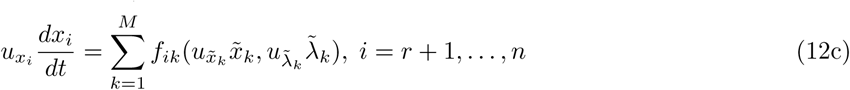

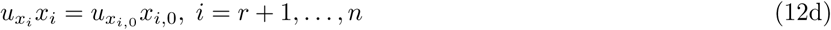

where *M* is the number of functional independent summands in the equation. It is convenient to rewrite Eq. (12c) as

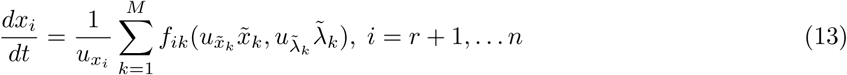

to perform the scale invariance analysis below in a simpler way.

If the solution is invariant under this transformation, then the right-hand sides of Eq. (10) and, consequently Eqs (12) should be equal. Besides, by the functional linear independence of the functions *f*_*ik*_ we can split each summand. Thus,

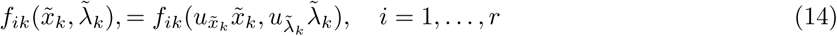

and

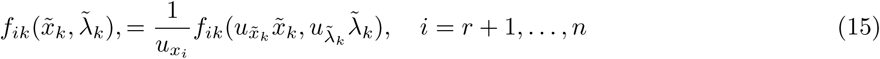

These new set of equations are much easier to solve than the ones that we would obtain from Eqs (12a)–(12c) (which would be equivalent to the so-called direct-test method [18]).

We can express the solution of these equations as

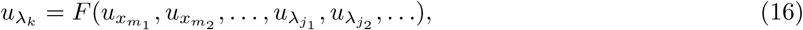

We denote these the **identifiability equations** of the model. For each parameter *k*, the identifiability equation will depend only on a few other scaling factors *m*_1_, *m*_2_, ….

If take the partial derivative of the transformed solution

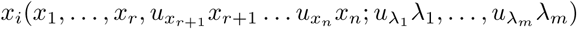

with respect to 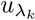, we find (by the chain rule)

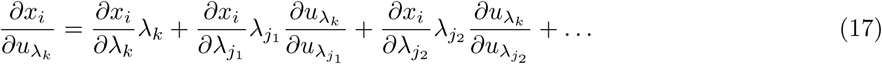

On the other hand, as the solution is invariant with respect to the scaling transformation, *x*_*i*_ should not depend on the arbitrary scaling factors, thus

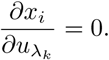

Finally, dividing Eq. (17) by *x*_*i*_ and defining

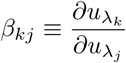

and the elasticity matrix *K* with elements *K*_*ij*_ given by

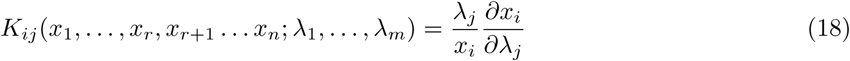

we find

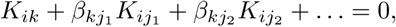

At this point we have two possibilities. Either all the coefficients β_*kj*_ = 0, meaning that Eq. (16) is not satisfied (we will simply have 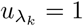), or the columns of the elasticity matrix related to the parameter λ_*k*_ has linearly dependent columns, and then the parameters λ_*k*_, λ_*j*_1, λ_*j*2_,… are unidentifiable. On the other hand, if the solution of the identifiability equation is 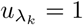 for some parameter, then the column of the elasticity matrix coming from that parameter is linearly independent of the others, and the parameter is **local** structurally identifiable. The adjective “local” is required because the method stems on the continuity of the derivative of the solution of Eq. (17). Thus, it is unable to capture any discrete (*countable*) transformations like, for instance, those related to exchanging two parameters of the model. Finally, by Eq. (12d) we find that 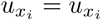, so an initial condition is identifiable whenever its corresponding variable is observable.

## Comparison with other methods

We have applied the method outlined in Box 1 to 13 different models defined and analyzed in detail in the Supporting Information. The choice is based on two criteria: on the one hand, models 1-5 are included for *pedagogical* purposes. They are simple enough to illustrate the method and most of the existing methods also provide definite answers. Models 6-13 were chosen because they have previously been analyzed using the methods summarized in the Introduction and in Table 1. This allows us to put our method in direct competition with those methods and to highlight their merits and limitations.

**Table 1.** List of current methods testing structural identifiability. We introduce here the acronyms referred to in Table 2.

| Method | Acronym | Main Ref. | Pros | Cons |
| --- | --- | --- | --- | --- |
| Direct test method | DT | [18, 20] | Simple | Limited |
| Implicit function theorem | IFT | [26] | Software | Limited |
| Taylor series approach | TS | [14] | Simple | Computationally Expensive |
| Generating series approach | GS | [13] | Simple, Software | Computationally Expensive |
| Similarity Transformation | ST | [16] | Software | Computationally Expensive |
| Differential algebra | DA | [29, 32, 34] | Software, Conclusive | Limited, Comp. Expensive |
| Reaction Network theory | RNT | [40, 41] | Simple, Hybrid with other | Only reaction systems |
| STRIKE-GOLDD | SG | [9, 22] | Powerful, Software | Computationally Expensive |
| Scaling Invariance Method | **SIM** | This work | Simple, Widely applicable | Only Local Identifiability |

The results of this comparison are summarized in Table 2, which is inspired by a similar table in Ref. [7].

**Table 2.** Summary of models compared in the literature: The number in brackets in the Model Name column correspond to the number of observed variables. Model Numbers correspond to those in Table 1 in the Supporting Information. The acronyms for the methods are summarized in Table 1. This table has been inspired by Table 1 in Ref. [7].

| Model name | Main Ref. | Model Number | Global Struct. Id. | Local Struct. Id. | Unidentifiable | Not Conclusive Not Applicable |
| --- | --- | --- | --- | --- | --- | --- |
| Goodwin model (1) | [7] | 6 |  |  | SG,**SIM** | TS,GS,ST,DT,DA,IFT,RNT |
| Goodwin model (all) | [7] | 6bis |  | TS,GS,IFT,RNT | DA,SG,**SIM** | ST,DT |
| Circadian clock model | [42] | 7 |  |  | TS,GS,RNT,SG,**SIM** | ST,DT,DA,IFT |
| HIV model (1) | [6, 43] | 8 |  |  | All |  |
| HIV model (2) | [6, 43] | 8bis | DA,IFT,RNT | TS,GS,**SIM** |  | DT,ST |
| Linear HIV model (1) | [6, 43, 44] | 8ter | DA,IFT,RNT,SG | DT,ST,TS,GS,**SIM** |  |  |
| Glycolysis model | [45] | 9 | GS,DA,RNT | TS,RNT,**SIM** |  | ST,DT |
| High dimensional model | [42] | 10 | TS,GS,DA,RNT | IFT,**SIM** |  | ST,DT |
| NF-κ model B (1) | [46] | 11 | | | SG, **SIM** | TS,GS,ST,DT,DA,IFT,RNT |
| NF-κ model B (2) | [46] | 11bis | GS,RNT | TS,**SIM** | SG | ST,DT,DA,IFT |
| Pharmacokinetics model (1) | [47] | 12 |  | TS,GS,RNT,SG,**SIM** |  | ST,DT,DA,IFT |
| Pharmacokinetics model (2) | [47] | 12bis | DA | GS,SG,**SIM** |  | ST,DT,IFT,RNT |
| Within-host virus model | [27] | 13 | DA | **SIM** |  | TS,GS,ST,DT,IFT,RNT |

## Discussion and Conclusions

Table 2 shows that our method can handle any complex model and provides a local structural identifiability criterion that is compatible with those methods capable of producing an answer. Thus, our method is widely applicable. It is worth noting that in several cases where our scaling method comes with a conclusive answer, other more complicated methods cannot address those cases (rightmost column in the table). As any global structural identifiable model is also local, our results are compatible with those methods that can address that difference.

It is worth emphasizing that we have performed our test by hand, as illustrated in the Supporting Information, and that, after some practice (and using some interesting *motifs* as having sums of different parameters, or the coefficients related to *diagonal* terms in the system of equations) the calculations can be made in a few minutes. This contrasts with the most sophisticated methods that, by hand, can fill several pages [27] or take hours using symbolic computation packages.

Together, this broad applicable and simplicity are the main features of our method and this may attract the interest of mathematical modelers and spread the *culture* of checking structural identifiability as a mandatory step when fitting experimental data.

We would like to highlight a connection with the so-called Buckingham-Π theorem of dimensional analysis [48]. In some sense, the scale invariance property is related to the principle of dimensional homogeneity, *i.e.*, the constraints on the functional form of the independent variables with the parameters. Our identifiability equations are therefore similar to finding the so-called Π-groups in the theorem. A limitation of the method is that it is restricted to testing local identifiability. This is implicit in the differentiability of the elasticity matrix which, by definition, is a local operation. Discrete symmetries are not captured, and more sophisticated methods (based on Lie group transformations [39]) are required. However, simple manipulation of the equations to remove the latent variables can improve the explanatory power of the method and might capture those discrete symmetries (see Sec. 3.8 of the Supporting Information). We leave that extension for future developments.

Finally, in this work we have chosen to solve the scaling factor equations directly as it is easy to perform with pen and paper. However, if we were to redefine the scaling factors as 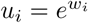, the new factors *w*_*i*_ would obey a linear system of homogeneous equations. It is therefore expected that the problem of identifiability is related to the rank of the matrix defining the linear system of equations. In that regard, the theorems presented in the Supporting Information could be supplemented with generic results on homogeneous systems of equations. Thus, our results provide a solid ground for the method and indicate a venue for further development in other systems like delay-differential or partial differential equations. Finally, while we emphasize the simplicity of the method, it is obviously amenable to be implemented using symbolic computation packages.

## Supporting information

In the Supporting Information we collect the theorems sustaining the method and a catalogue of models with a detailed computation of the identifiability equations that were used to build Table 2.

## Acknowledgments

This work has been partially supported by grant FIS2016-78883-C2-2-P (Ministerio de Economia, Industria y Competitividad - Agencia Estatal de Investigacion). This work was initiated during summer visits of the authors to the Los Alamos National Laboratory, and we thank Nick Hengartner and Alan Perelson (LANL) for their hospitality and helpful comments on this work, and the Santa Fe Institute for supporting the summer visits of RdB.

