## Supporting Information for "Testing structural identifiability by a simple scaling method"

**a** Grupo Interdisciplinar de Sistemas Complejos (GISC)
**b** Instituto de Investigación Tecnológica (IIT), Universidad Pontificia Comillas, Madrid, E28015, Spain.
**c** Theoretical Biology and Bioinformatics, Utrecht University, Utrecht, The Netherlands.

‡All authors also contributed equally to this work.
*

**3 Summary Table** **13**

### 1 Functional independence theorems

In this section we introduce some general results necessary to prove our main results throughout the paper

#### 1.1 The Wronskian theorem

The Wronskian theorem (see, *e.g*, Ref. [11]) states that, if $n$ functions $f_1(x) \ldots f_n(x)$ are *linearly independent*, then any of the following statements hold:

1. The only solution of the equation
$$\sum_{i=1}^{n} a_i f_i(x) = 0 \tag{1}$$
is $a_1 = \cdots = a_n = 0$.

2. The Wronskian, $W$, satisfies
$$W = \det \begin{pmatrix} f_1(x) & f_2(x) & \dots & f_n(x) \\ f_1'(x) & f_2'(x) & \dots & f_n'(x) \\ \dots & \dots & \dots & \dots \\ f_1^{(n-1)} & f_2^{(n-1)} & \dots & f_n^{(n-1)} \end{pmatrix} \neq 0 \tag{2}$$

For example, if $f_1(x) \equiv f(x)$, $f_2(x) = 1$, then Eq. (1) is
$$a_1 f(x) + a_2 = 0 \tag{3}$$
and, as $W = -f'(x) \neq 0$ (except for the trivial case $f(x)$ =constant) then $a_1 = a_2 = 0$.

##### 1.2 A general result for functional equations in many variables

Consider the general functional equation
$$f_1(x_1) = f(x_2) \tag{4}$$
where $x_1$ and $x_2$ are independent variables. Taking the partial derivative with respect to $x_2$ (alternatively, $x_1$) then
$$\frac{df_2}{dx_2} = 0 \Rightarrow f_2(x_2) = K \text{ (a constant)},$$
hence
$$f(x_1) = K$$
that has the same form as Eq. (3) (if we identify $a_2 = -K$) so $K = 0$ and, hence
$$f_1(x_1) = f_2(x_2) = 0. \tag{5}$$
In general, the only solution of the equation
$$\sum_{i=1}^{n} f_i(x_i) = 0 \tag{6}$$
is
$$f_1(x_1) = f_2(x_2) = \cdots = f_n(x_n) = 0. \tag{7}$$
This is an example of the method of separation of variables, that is a particular case of the generalized Wronskian theorem of the next section.

##### 1.3 Generalized Wronskian

**Theorem (from Ref. [12]):** Let $\Delta_0, \dots \Delta_{n-1}$ be differential operators that are combinations of terms of the form
$$\Delta_s = \left(\frac{\partial}{\partial x_1}\right)^{j_1} \cdots \left(\frac{\partial}{\partial x_m}\right)^{j_m}, \text{ with } j_1 + \ldots + j_m \leq s,$$
then the if the generalized Wronskian, $\mathcal{W}$, over the functions $f_i(x_1, \ldots, x_m)$, with $i = 1 \ldots n$,
$$\mathcal{W} = \begin{pmatrix} \Delta_0 f_1 & \cdots & \Delta_0 f_n \\ \cdots & \cdots & \cdots \\ \Delta_{n-1} f_1 & \cdots & \Delta_{n-1} f_n \end{pmatrix} \tag{8}$$
is different from zero (except in isolated points) then those functions $f_i$ are linearly independent.

#### 1.4 Some examples of linearly independent functions

Let us see some examples deriving from the previous results.

##### 1.4.1 Polynomial equations

Consider the equation

$$a_0 + a_1 x + a_2 x^2 = 0 \tag{9}$$

If we define $f_i(x) = x^i$ then the (traditional) Wronskian is given by

$$W = \det \begin{pmatrix} 1 & x & x^2 \\ 0 & 1 & 2x \\ 0 & 0 & 2 \end{pmatrix} = 2 \neq 0,$$

and 1, $x$ and $x^2$ are linearly independent functions and the solution of Eq. (9) is $a_0 = a_1 = a_2 = 0$.

##### 1.4.2 Polynomial-rational equations

Let us now consider the equation

$$a_1 x + b_1 \frac{x}{1+x} = 0. \tag{10}$$

If we define $f_1 = x$ and $f_2 = \frac{x}{1+x}$ then the Wronskian is given by

$$W = \det \begin{pmatrix} x & \frac{x}{1+x} \\ 1 & \frac{1}{(x+1)^2} \end{pmatrix} = -\frac{x^2}{(x+1)^2} \neq 0 \tag{11}$$

for generic values of $x \neq 0$, so the functions $f_1$ and $f_2$ are linearly independent and, consequently, the solution of Eq. (10) is $a_1 = b_1 = 0$.

##### 1.4.3 Examples with multiple variables

Consider again Eq. (4), in this case $n = 2$ (two functions) and $m = 2$ (two variables), then

$$\Delta_0 = 1 \qquad \Delta_1 = \frac{\partial}{\partial x_1} + \frac{\partial}{\partial x_2}$$

and the generalized Wronskian is

$$\mathcal{W} = \begin{pmatrix} f_1(x_1) & f_2(x_2) \\ f_1'(x_1) & f_2'(x_2) \end{pmatrix} = f_1(x_1) f_2'(x_2) - f_2(x_2) f_1'(x_1) \neq 0$$

for generic values of $x_1$ and $x_2$, proving Eq. (7) by different means.
More interesting is the case in which $f_1$ and $f_2$ can depend on both $x_1$ and $x_2$. For instance, consider the following equation:

$$a_1 \frac{1}{1+x_1} + a_2 x_1 x_2 = 0.$$

After some manipulation, it can be written as Eq. (4), with $f_1(x_1) = 1/x_1$ and $f_2(x_2) = x_2$, namely

$$a_1 \frac{1}{x_1} + a_2 x_2 = 0.$$

However, it is illustrative to prove the same result using the generalized Wronskian theorem, Eq. (8). Thus,

$$\mathcal{W} = \begin{pmatrix} \frac{1}{1+x_1} & x_1 x_2 \\ -\frac{1}{(1+x_1)^2} & x_1 + x_2 \end{pmatrix} = \frac{x_1^2 + 2x_1 x_2 + x_1 + x_2}{(x_1+1)^2} \neq 0$$

consequently, $a_1 = a_2 = 0$.

### 2 A catalogue of models

#### 2.1 Model 1: A simple one-dimensional linear model

Consider a simple *exponential* model given by

$$\dot{x} = abx \tag{12}$$

where $\dot{x}$ denotes the time derivative of the variable $x$ and $a$, $b$ are two (constant) parameters.

##### Identifiability equations

In order to apply our method, we make the following substitutions:

$$\begin{cases} a & \rightarrow & u_a a \\ b & \rightarrow & u_b b \end{cases} \tag{13}$$

Because the variable $x$ is observed in the experiment, we find $u_x = 1$. Substituting into Eq. (12) gives

$$\dot{x} = u_a u_b abx.$$

If the system is identifiable, the scaled model should be invariant under such a transformation (hereafter we will refer to these equations as the **invariance conditions**). Thus,

$$u_a u_b abx = abx$$

For values of $a$, $b$ and $x$ different from zero, the solution of that equation is

$$\{u_a u_b = 1 \text{ or}, \; u_b = \frac{1}{u_a}, \tag{14}$$

which implies that there are an *infinite* number of transformations of the form Eq. (13) that would provide the same experimental data and, so the system of Eq. (12) is unidentifiable (as at least one parameter is). Equation (14) also predicts that $ab$ is an identifiable group of parameters.
In this simple case, this result could have been found by simple inspection of the exact solution. Thus,

$$x(t) = x(0)e^{abt},$$

where clearly one can see that multiplying $a$ by an arbitrary constant $u_a$ and dividing $b$ by $1/u_a$ as in Eq. (14).
Another simple case is

$$\dot{x} = (a - b)x. \tag{15}$$

The transformation of Eq. (13) leads to the invariance condition,

$$(u_a a - u_b b) = (a - b)$$

meaning that there are an infinite number of solutions corresponding to all the values of $u_{a,b}$ lying in the line

$$u_a = 1 - \frac{b}{a}(1 - u_b). \tag{16}$$

It is worth noting that Eq. (16) is a direct reflection of the more obvious (*shift*) symmetry of Eq. (15),

$$\begin{cases} a \rightarrow a + \Delta \\ b \rightarrow b + \Delta \end{cases},$$

so

$$\dot{x} = ((a + \not{\Delta}) - (b + \not{\Delta}))x = (a - b)x.$$

#### 2.2 Model 2: Unidentifiable nonlinear model [1]

Our second example was introduced in Ref. [1] as an unidentifiable nonlinear model with two variables, when only $x_2$ is observed. In particular,

$$\dot{x}_1 = p_1 x_1^2 + p_2 x_1 x_2 + g, \tag{17a}$$
$$\dot{x}_2 = p_3 x_1^2 + p_4 x_1 x_2, \tag{17b}$$
$$x_1(0) = 0, \tag{17c}$$
$$x_2(0) = 0, \tag{17d}$$
$$x_1 \text{ is observed} \tag{17e}$$
$$g \text{ is a known input function.} \tag{17f}$$

In this work we denote input functions as $g(t)$ instead of the usual convention $u(t)$ to avoid the confusion with the scaling factors, $u_\lambda$.

**Identifiability equations**

In this case,

$$\begin{cases} x_2 & \to & u_{x_2} x_2 \\ p_1 & \to & u_{p_1} p_1 \\ p_2 & \to & u_{p_2} p_2 \\ p_3 & \to & u_{p_3} p_3 \\ p_4 & \to & u_{p_4} p_4 \end{cases}, \tag{18}$$

as $x_1$ is observed (so, $u_{x_1}$)1). We can define the functional linear independent functions:

$$f_{11} = p_1 x_1^2 \qquad f_{12} = p_2 x_1 x_2 \qquad f_{21} = p_3 x_1^2 \qquad f_{22} = p_4 x_1 x_2,$$

and from Eqs [14]-[15] in the main text

$$u_{p_1} p_1 x_1^2 = p_1 x_1^2 \qquad u_{p_2} u_{x_2} p_2 x_1 x_2 = p_2 x_1 x_2$$

and

$$\frac{u_{p_3}}{u_{x_2}} p_3 x_1^2 = p_3 x_1^2 \qquad u_{p_4} p_4 x_1 x_2 = p_4 x_1 x_2$$

respectively. Hence, the identifiability equations are

$$\begin{cases} u_{p_1} = 1 \\ u_{p_2} u_{x_2} = 1 \\ u_{p_3} = u_{x_2} \\ u_{p_4} = 1 \end{cases} \tag{19}$$

As the system has more than 1 solution besides the trivial ($u_{p_1} = u_{p_2} = \ldots = 1$) it follows that the model is unidentifiable. Moreover, as discussed in the main text, Eq. (19) allows one to conclude that (i) if $x_2$ were to be observed ($u_{x_2} = 1$), all the parameters would be identifiable, and (ii) the combination $u_{p_2} u_{p_3}$ is identifiable as, for any *scale* of $x_2$, the condition $u_{p_2} u_{p_3} = 1$ is always fulfilled and hence $p_2 p_3$ is an identifiable group (as concluded in Ref. [1]).

#### 2.3 Model 3: A local structurally identifiable nonlinear model [1]

In Ref. [1] the authors also present a nonlinear model that is locally identifiable, *i.e.*,

$$\dot{x}_1 = p_1 x_1 x_2 + g, \tag{20a}$$
$$\dot{x}_2 = p_2 x_1 x_2 + p_3 x_2^2, \tag{20b}$$
$$x_1(0) = p_4, \tag{20c}$$
$$x_2 \text{ is observed} \tag{20d}$$
$$g \text{ is a known input function.} \tag{20e}$$

###### Identifiability equations

In this case,

$$\begin{cases} x_1 & \to & u_{x_1}x_1 \\ p_1 & \to & u_{p_1}p_1 \\ p_2 & \to & u_{p_2}p_2 \\ p_3 & \to & u_{p_3}p_3 \end{cases}, \tag{21}$$

and, from the initial condition Eq. (20c), $u_{p_4} = u_{x_4}$.
Following the same arguments as in the preceding example (the functions $f_{ik}$ are the same), we conclude that the identifiability equations are

$$\begin{cases} u_{p_1} = 1 \\ u_{p_2}u_{x_1} = 1 \\ u_{p_3} = 1 \\ u_{p_4}/u_{x_1} = 1 \\ u_{x_1} = 1 \end{cases} \tag{22}$$

where the last equation is a consequence of the fact that $u$ in Eq. (20a) is a known function (in the jargon of control theory, it is an *input*). The only solution is the trivial one, $u_{p_1} = \ldots = u_{p_4} = u_{x_1} = 1$, so the system is local structurally identifiable.

#### 2.4 Model 4: A Lotka-Volterra-like model

Let us consider now the following system,

$$\dot{x}_1 = rx_1(1 - x_1/K) - ax_1x_2, \tag{23a}$$

$$\dot{x}_2 = cax_1x_2 - dx_2 - ex_2^2, \tag{23b}$$

$$\dot{x}_3 = px_2 - fx_3. \tag{23c}$$

###### Identifiability equations

Again, the linear independent functions $f_{ik}$ are polynomials of the variables $x_1$, $x_2$ and $x_3$. So, the identifiability equations are

$$\begin{cases} \left.\begin{aligned} u_r &= 1, \\ u_r u_{x_1} &= u_K, \\ u_a u_{x_2} &= 1, \end{aligned}\right\} \text{From Eq. } (23a) \\ \left.\begin{aligned} u_c u_a u_{x_1} &= 1, \\ u_d &= 1, \\ u_e u_{x_2} &= 1, \end{aligned}\right\} \text{From Eq. } (23b) \\ \left.\begin{aligned} u_p u_{x_2} &= u_{x_3}, \\ u_f &= 1. \end{aligned}\right\} \text{From Eq. } (23c) \end{cases} \tag{24}$$

so, $r$, $d$ and $f$ are always identifiable. In general we need to consider different cases:

- *Case 1: Only $x_3$ is observed in the experiment ($u_{x_3} = 1$)*

  With this information, the identifiable parameters would be

  $$r, x_1(0)/K, ax_2(0), cax_1(0), d, ex_2(0), px_2(0), f.$$

  meaning that the model is unidentifiable. Here $x_1(0)$ and $x_2(0)$ are the *unknown* parameters for the initial conditions. As revealed in the next example, the conclusion that only $x_1(0) = u_K$ is a direct consequence of the theory and the fact that their ratio is dimensionless in Eq. (23a).

- *Case 2: $x_3$ and $x_1$ are observed in the experiment ($u_{x_3} = u_{x_1} = 1$)*

  Now, the identifiable groups are:

  $$r, K, ax_2(0), ca, d, ex_2(0), px_2(0), f,$$

  so the model remains unidentifiable.

- *Case 3: $x_3$ and $x_2$ are observed in the experiment ($u_{x_3} = u_{x_2} = 1$)*

  Now, the identifiable groups are:

  $$r, x_1(0)/K, a, cx_1(0), d, e, p, f,$$

  so the model is unidentifiable.

- *Case 4: $x_1$, $x_2$ and $x_3$ are observed in the experiment*

  In this case, the model is local structurally identifiable.

#### 2.5 Model 5: A batch reactor (Michalis-Menten-like) model [2]

This example is of special interest in Theoretical Biology for its ubiquity in enzymatic reactions.

$$\dot{x} = \frac{\mu s x}{K_s + s} - K_d x, \tag{25a}$$

$$\dot{s} = \frac{-\mu s x}{Y(K_s + s)}, \tag{25b}$$

$$x(0) = b_1, \tag{25c}$$

$$s(0) = b_2, \tag{25d}$$

$$x \text{ is observed} \tag{25e}$$

###### Identifiability equations

In this case, we identify the following linearly independent functions from the first equation

$$f_{x1}(s) = \frac{\mu s x}{K_s + s} \qquad f_{x2}(x) = -K_d x$$

with generalized Wronskian

$$\frac{K_d K_s \mu x^2}{(K_s + s)^2} \neq 0 \text{ for generic values of } x, s.$$

Hence, the invariant equations are simply

$$\begin{cases} \frac{\mu}{K_s+s} = \frac{u_\mu u_s \mu}{u_{K_s} K_s + u_s s} & \rightarrow \qquad K_s + s = \frac{u_{K_s} K_s + u_s s}{u_\mu u_s} \\ K_d = u_{K_d} K_d & \\ \frac{\mu}{Y(K_s+s)} = \frac{u_\mu \mu}{u_Y Y(u_{K_s} K_s + u_s s)} & \rightarrow \quad Y(K_s + s) = \frac{u_Y Y(u_{K_s} K_s + u_s s)}{u_\mu} \end{cases},$$

and the identifiability equations are,

$$\begin{cases} u_\mu u_s = 1, \\ u_{K_s} = 1, \\ u_{K_d} = 1, \\ u_{K_s} = u_s, \\ u_\mu = 1, \\ u_Y u_{K_s} = 1, \\ u_Y u_s = 1, \end{cases} \tag{26}$$

From the initial conditions, $u_{b_1} = 1$ and $u_{b_2} = u_s$. Because of the fourth and the last two equations, we cannot identify uniquely all the parameters and, hence, the model is unidentifiable. It is worth noting that using the Taylor series approach for this model required several pages of involved calculations [13]. An interesting corollary of this example illustrates an important consequence of linear functional independence and dimensional analysis. In particular, by solving the system of equations we arrive at the conclusion that $u_{K_s} = u_s$. This is independent of the mathematical form of the numerator and simplifies qualitatively our identifiability analysis. This dimensional argument also implies that if have an equation of the form

$$\dot{x}_i = p f(\tilde{x}, \tilde{\lambda}) \tag{27}$$

and $f(\cdot)$ is **dimensionless** function, then it follows that $u_{x_i} = u_p$.

#### 2.6 Model 6: Goodwin's model [3]

In contrast with Model 5, this model contains Hill functions with unknown exponent. Specifically,

$$\dot{x}_1 = -bx_1 + \frac{a}{A + x_3^\sigma}, \tag{28a}$$

$$\dot{x}_2 = \alpha x_1 - \beta x_2, \tag{28b}$$

$$\dot{x}_3 = \gamma x_2 - \delta x_3, \tag{28c}$$

$$\tag{28d}$$

As in Ref. [3] we will consider two scenarios: (i) only $x_1$ is observed and (2) all variables are observed.

**Identifiability equations**

Using the generalized Wronskian, it is easy to see that $f_{11}(x_1) = bx_1$, $f_{12}(x_2) = \frac{a}{A+x_3^\sigma}$ are linearly independent functions. This can be proven easily as, for instance,

$$\mathcal{W} = \det \begin{pmatrix} \frac{a}{x_3^\sigma + A} & x_3 \\ -\frac{a x_3^{\sigma-1}\sigma}{\left(x_3^\sigma + A\right)^2} & 1 \end{pmatrix} = \frac{a\left(A + (\sigma+1)x_3^\sigma\right)}{\left(A + x_3^\sigma\right)^2} \neq 0$$

for generic values of the parameters. Thus, the identifiability equations are (assuming that only $x_1$ is observed)

$$\begin{cases} u_b = 1 \\ u_A = u_a \\ u_\sigma = 1 \\ u_{x_3}^\sigma = u_a \\ u_\beta = 1 \\ u_\alpha = u_{x_2} \\ u_\delta = 1 \\ u_\gamma u_{x_2} = u_{x_3} \end{cases},$$

meaning that the system is unidentifiable. However, if $x_2$ and $x_3$ are observed, then $u_{x_2} = u_{x_3} = 1$ and the system is local structurally identifiable. Interestingly, our method does not require the transformation of the original problem into a polynomial one as was done in Ref. [3]. Like in the Michalis-Menten example, combining Eqs. $u_A = u_a$ and $u_{x_3}^\sigma = u_a$, allows us to conclude that in this case the dimensions of $A$ and $x_3$ are constrained which, again, simplifies the analysis.

#### 2.7 Model 7: Circadian clock [3, 4]

This model has 7 variables and 28 parameters following the equations,

$$\dot{x}_1 = n_1 \frac{x_6^a}{g_1^a + x_6^a} - m_1 \frac{x_1}{k_1 + x_1} + q_1 x_7 g(t), \tag{29a}$$

$$\dot{x}_2 = p_1 x_1 - r_1 x_2 + r_2 x_3 - m_2 \frac{x_2}{k_2 + x_2}, \tag{29b}$$

$$\dot{x}_3 = r_1 x_2 - r_2 x_3 - m_3 \frac{x_3}{k_3 + x_3}, \tag{29c}$$

$$\dot{x}_4 = n_2 \frac{g_2^2}{g_2^2 + x_3^2} - m_4 \frac{x_4}{k_4 + x_4}, \tag{29d}$$

$$\dot{x}_5 = p_2 x_4 - r_3 x_5 + r_4 x_6 - m_5 \frac{x_5}{k_5 + x_5}, \tag{29e}$$

$$\dot{x}_6 = r_3 x_5 - r_4 x_6 - m_6 \frac{x_6}{k_6 + x_6}, \tag{29f}$$

$$\dot{x}_7 = p_3 - m_7 \frac{x_7}{k_7 + x_7} - (p_3 + q_2 x_7) g(t), \tag{29g}$$

$$g(t) \text{ is a known input function}, \tag{29h}$$

$$x_i(0) = 0 \text{ (for } i = 1, \ldots, 7), \tag{29i}$$

$$x_1, x_4 \text{ are observed} \tag{29j}$$

**Identifiability equations**

In this case, all the terms on the right hand side of Eqs. (29) are of the same form as in previous models, so they are all linearly independent functions. Thus, we can obtain the identifiability equations by equating each term to itself after the transformations $x_k \to u_{x_k} x_k$, etc ... (and, because of Eq. (29j), $u_{x_1} = u_{x_4} = 1$). In this sort of problems, it is not worth it writing down all the identifiability equations. Instead, it is easier to perform the analysis progressively. The analyses of previous models have shown us that the rate terms (namely, a linear term proportional to $x_k$ in the equation of $\dot{x}_k$) are always locally identifiable. In this problem, this implies that $u_{r_1} = \ldots = u_{r_4} = 1$, and using this, $u_{x_3} = u_{x_2}$ and $u_{x_6} = u_{x_5}$. Similarly, from the dimensional arguments in models 5 and 6 and Eq. (27), $k_1 \ldots k_7$ and $m_1 \ldots m_7$ have the same scaling factors as $x_1 \ldots x_7$, respectively. Because $x_1$ and $x_4$ are observed, $u_{m_1} = u_{k_1} = u_{m_4} = u_{k_4} = 1$. Analogously, $u_{n_1} = u_{n_2} = 1$. After solving the first equations it follows that the parameters $k_i$ (for $i \neq 1, 4$) cannot be identified, and that the system is unidentifiable. This iterative way of solving helps to simplify the identifiability equations and reduces the complexity into a *back-of-the-envelope* analysis.

#### 2.8 Model 8: the HIV model with an eclipse phase [5, 6]

The virus dynamics model with an additional *compartment* ($I_2$ that accounts for the so-called eclipse phase

$$\dot{T} = -\beta T V, \tag{30a}$$

$$\dot{I}_1 = \beta T V - k I_1, \tag{30b}$$

$$\dot{I}_2 = k I_1 - \delta I_2, \tag{30c}$$

$$\dot{V} = p I_2 - c V, \tag{30d}$$

$$T(0) = T_0, \tag{30e}$$

$$I_1(0) = I_{1,0}, \tag{30f}$$

$$I_2(0) = I_{2,0}, \tag{30g}$$

$$V(0) = V_0. \tag{30h}$$

In this case,

$$\begin{aligned} f_{11} &= f_{21} = \beta T V, \\ f_{22} &= f_{31} = k I_1, \\ f_{32} &= f_{41} = \delta I_2, \\ f_{42} &= c V. \end{aligned}$$

so $f_{ij}$ and $f_{ik}$ are linearly independent functions for all $j \neq k$.
First, we will consider the typical case where only $V$ is observed.

$$\begin{cases} u_\beta = 1 \\ u_\beta u_T = u_{I_1} \\ u_k = 1 \\ u_k u_{I_1} = u_{I_2} \\ u_\delta = 1 \\ u_p u_{I_2} = 1 \\ u_c = 1 \end{cases}$$

making the system is unidentifiable. However, if any of the other variables are observed ($T$, $I_1$ or $I_2$) then the system becomes locally structurally identifiable as the only solution would be all the scaling factors equal to 1. A well-known interesting case appears when the eclipse phase is neglected (*i.e.*, $I_1$ is dropped) and we consider a treatment where novel infections are totally inhibited ($\beta = 0$) [14]. In that case, the system reduces to two-dimensional linear system of ODEs

$$\dot{I}_2 = -\delta I_2, \tag{31a}$$

$$\dot{V} = p I_2 - c V, \tag{31b}$$

$$I_2(0) = I_{2,0}, \tag{31c}$$

$$V(0) = V_0. \tag{31d}$$

Using our scaling invariance method one would conclude that only $V_0$, $c$ and $\delta$ and the group $pI_{2,0}$ are local structurally identifiable (when $V$ is observed). However, as the model is linear the exact solution is

$$I_2(t) = \frac{cV_0}{p} e^{-\delta t}, \qquad \text{and} \qquad V(t) = \frac{V_0}{c-\delta}\left(ce^{-\delta t} - de^{-ct}\right).$$

This confirms that, as $I_2$ is not observed, that the product $pI_{2,0}$ is identifiable as a group, and that $c$ and $d$ are exchangeable in the solution for $V(t)$. Thus one cannot uniquely determine them. This is related to the limitation of our model to test local structural identifiability only: these parameters can be estimated only in a subset of the space of parameters.
In this particular case, the method recovers its predictive ability if Eqs. (31) are manipulated to eliminate the latent variable, $I$. In particular, taking the time derivative in Eq. (31b), and use Eq. (31a), then

$$\frac{d^2V}{dt^2} = p\frac{dI_2}{dt} - c\frac{dV}{dt} \Rightarrow \frac{dV^2}{dt^2} = -c\delta V - (c+\delta)\frac{dV}{dt},$$

and our method would provide the identifiability equations

$$\begin{cases} u_c u_\delta &= 1 \\ u_c c + u_\delta \delta &= c + \delta \end{cases}$$

which has only two solutions (*a finite number*)

$$\{u_c \to 1, u_d \to 1\ , \qquad \left\{u_c \to \frac{\delta}{c}, u_d \to \frac{c}{\delta}\right.$$

explaining why they are locally but not globally identifiable.

##### 2.9 Model 9: Glycolysis inspired metabolic pathway [7]

$$\dot{x}_1 = -\frac{k_1 x_1}{x_1 + k_M} g_1, \tag{32a}$$

$$\dot{x}_2 = \frac{k_1 x_1}{x_1 + k_M} g_1 - \frac{k_2 x_2}{x_2 + k_M} g_2, \tag{32b}$$

$$\dot{x}_3 = \frac{k_2}{x_2 + k_M} g_2 - \frac{k_3 x_3}{x_3 + k_M} g_3, \tag{32c}$$

$$\dot{x}_4 = \frac{k_2 x_2}{x_2 + k_M} g_2 + \frac{k_3 x_3}{x_3 + k_M} g_3 - \frac{k_4 x_4}{x_4 + k_M} g_4 \tag{32d}$$

$$\dot{x}_5 = \frac{k_4 x_4}{x_4 + k_M} g_4, \tag{32e}$$

$$x_i(0) = S_i, \quad i = 1 \ldots 5, \tag{32f}$$

$$\text{All states are observed,} \tag{32g}$$

and $g_1 \ldots g_4$ are known dimensionless input functions. As all the states are observed, so they are all the initial conditions, $S_i$. Besides, from the dimensional analysis discussion in previous examples, we can readily conclude that all the parameters are identifiable: $k_M$, because it has the same dimensions as all the $x$'s, and $k_1 \ldots k_4$ because the functions

$$\frac{x_i g_i}{x_i + K_M}$$

are dimensionless. So the system is locally structurally identifiable.

##### 2.10 Model 10: A high dimensional non-linear model [3, 4]

$$\dot{x}_1 = -v_{max}\frac{x_1}{k_m + x_1} - p_1 x_1 + g(t), \tag{33a}$$

$$\dot{x}_i = p_{i-1}x_{i-1} - p_i x_i, \qquad i = 2 \ldots 20, \tag{33b}$$

$$g(t) \text{ is a known input function.} \tag{33c}$$

$$\text{All states are observed} \tag{33d}$$

Despite the dimensionality of the problem, the fact that all the equations contain a *diagonal* (rate) term of the form $-p_i x_i$, implies that $u_{p_1} = \cdots = u_{p_{20}} = 1$. Finally, from the first equation $u_{k_m} = u_{v_{max}} = 1$, and hence the problem is local structurally identifiable.

#### 2.11 Model 11: NF-$\kappa$B signaling pathway [3, 8]

This is an inhibitor/activator model introduced in Ref. [4, 8] with 15 variables and 29 parameters.

$$
\begin{aligned}
\dot{x}_1 &= k_{prod} - k_{deg}x_1 - k_1 x_1 g(t), && \text{(34a)}\\
\dot{x}_2 &= -(k_3 + k_{deg})x_2 - a_2 x_2 x_{10} + t_1 x_4 - a_3 x_2 x_{13} + t_2 x_5 + (k_1 x_1 - k_2 x_2 x_8)g(t), && \text{(34b)}\\
\dot{x}_3 &= k_3 x_2 - k_{deg}x_3 + k_2 x_2 x_8 g(t), && \text{(34c)}\\
\dot{x}_4 &= a_2 x_2 x_{10} - t_1 x_4, && \text{(34d)}\\
\dot{x}_5 &= a_3 x_2 x_{13} - t_2 x_5, && \text{(34e)}\\
\dot{x}_6 &= c_{6a} x_{13} - a_1 x_6 x_{10} + t_2 x_5 - i_1 x_6, && \text{(34f)}\\
\dot{x}_7 &= i_1 kv x_6 - a_1 x_{11} x_7, && \text{(34g)}\\
\dot{x}_8 &= c_4 x_9 - c_5 x_8, && \text{(34h)}\\
\dot{x}_9 &= c_2 + c_1 x_7 - c_3 x_9, && \text{(34i)}\\
\dot{x}_{10} &= -a_2 x_2 x_{10} - a_1 x_{10} x_6 + c_{4a} x_{12} - c_{5a} x_{10} - i_{1a} x_{10} + e_{1a} x_{11}, && \text{(34j)}\\
\dot{x}_{11} &= -a_1 x_{11} x_6 - c_{6a} x_{13} - a_3 x_2 x_{13} + e_{2a} x_{14}, && \text{(34k)}\\
\dot{x}_{12} &= c_{2a} + c_{1a} x_7 - c_{3a} x_{12}, && \text{(34l)}\\
\dot{x}_{13} &= a_1 x_{10} x_6 - c_{6a} x_{13} - a_3 x_2 x_{13} + e_{2a} x_{14}, && \text{(34m)}\\
\dot{x}_{14} &= a_1 x_{11} x_7 - e_{2a} kv x_{14}, && \text{(34n)}\\
\dot{x}_{15} &= c_{2c} + c_{1c} x_7 - c_{3c} x_{15}, && \text{(34o)}\\
g(t) &\text{ is a known input function,} && \text{(34p)}\\
&x_2, x_7, x_9, x_{12} \text{ are observed,} && \text{(34q)}\\
&x_{10} + x_{13}, x_1 + x_2 + x_3 \text{ are observed.} && \text{(34r)}
\end{aligned}
$$

##### Identifiability equations

Like Model 7, this problem can be solved iteratively. However, this deserves special attention as Eq. (34r) couples the observations of different variables (for instance, only the combination $x_{10} + x_{13}$ is observed). We are going to split this model in two cases:

- *Case 1: All the parameters are unknown.* This case is particularly simple to analyze as $x_{15}$, $c_{2c}$ and $c_{1c}$ only appear in Eq. (34o) so the identifiability equations coming from that are

$$
\begin{cases} u_{c_{2c}} = u_{x_{15}} \\ u_{c_{1c}} = u_{x_{15}} \end{cases},
$$

  meaning that the problem is unidentifiable. Of course a more thorough analysis to determine identifiable groups deserves further effort as in Case 2.

- *Case 2: Only $t_1, t_2, c_{3a}, c_{4a}, c_5, k_1, k_2, k_4, k_{prod}, k_{deg}, i_1, e_{2a}, i_{1a}$ are unknown, as in Ref. [3].* Since $c_{2c}$ and $c_{1e}$ are known from the literature, $x_{15}$ is observable. Then, iteratively we can unveil the identifiable (and observable) parameters. For instance, from the linear (rates) terms, we identify $k_{deg}$, $t_1$, $t_2$, $i_1$, .... From those and the observable variables, we identify $k_1$, $k_3$, and so on. We finally conclude that, if all the parameters except $t_1, t_2, c_{3a}, c_{4a}, c_5, k_1, k_2, k_4, k_{prod}, k_{deg}, i_1, e_{2a}, i_{1a}$ are known, then the model is locally structurally identifiable.

#### 2.12 Model 12: Pharmacokinetics model [9]

Following the formulation in Ref. [3],

$$\dot{x}_1 = \alpha_1(x_2 - x_1) - \frac{k_a v_m x_1}{k_c k_a + k_c x_3 + k_a x_1}, \tag{35a}$$
$$\dot{x}_2 = \alpha_2(x_1 - x_2), \tag{35b}$$
$$\dot{x}_3 = \beta_1(x_4 - x_3) - \frac{k_c v_m x_3}{k_c k_a + k_c x_3 + k_a x_1}, \tag{35c}$$
$$\dot{x}_4 = \beta_2(x_3 - x_4), \tag{35d}$$
$$x_1(0) = c_0, \tag{35e}$$
$$x_2(0) = 0, \tag{35f}$$
$$x_3(0) = \gamma c_0, \tag{35g}$$
$$x_4(0) = 0, \tag{35h}$$
$$x_1 \text{ and/or } x_2, \text{ are observed.} \tag{35i}$$

**Identifiability equations**

In this case,

$$\begin{cases} u_{\alpha_1} = 1 \\ u_{\alpha_1} u_{x_2} = 1 \\ u_{k_a} u_{v_m} = u_{k_a} \\ u_{k_a} u_{v_m} = u_{k_c} u_{k_a} \\ u_{k_a} u_{v_m} = u_{k_c} u_{x_3} \\ u_{\alpha_2} = 1 \\ u_{\alpha_2} = u_{x_2} \\ u_{\beta_1} = 1 \\ u_{\beta_1} u_{x_3} = u_{x_4} \\ u_{k_c} u_{v_m} = u_{k_a} \\ u_{k_c} u_{v_m} = u_{k_c} u_{k_a} \\ u_{k_c} u_{v_m} = u_{k_c} u_{x_3} \\ u_{\beta_2} = 1 \\ u_{\beta_2} u_{x_3} = u_{x_4} \\ u_{c_0} = 1 \\ u_{x_3} = u_\gamma u_{c_0}. \end{cases}$$

The unique solution is that all scaling factors, $u$, are 1 so the system is locally structurally identifiable. Again, we can arrive at the same conclusion more readily by analyzing the dimensions and the diagonal terms of the equation. From the first equation, we can rewrite the second term as

$$v_m \frac{x_1}{k_c + (k_c/k_a)x_3 + x_1}$$

so $u_{k_c} = u_{x_1}$, $u_{v_m} = 1$, $u_{k_c} = 1$ and $u_{x_3} = 1$. From the linear (rates) terms, we find $u_{\alpha_1} = u_{\alpha_2} = u_{\beta_1} = u_{\beta_2} = 1$ and, consequently $u_{x_3} = u_{x_4} = 1$. Finally, from the initial conditions, $u_{c_0} = 1$ and from all the above, $u_\gamma = 1$.

#### 2.13 Model 13: Within-host infectious disease [10]

Our final example is a non-linear model derived in the context of within-host infectious diseases. The non-linearity is neither polynomial or Hill-type.

$$\dot{H} = -\alpha P, \tag{36a}$$
$$\dot{V} = \lambda P - \eta V - \beta H V, \tag{36b}$$
$$\dot{P} = \beta H V - \alpha P - \alpha\gamma \frac{P^2}{H}, \tag{36c}$$
$$V \text{ is observed,} \tag{36d}$$

**Identifiability equations**

In this case,

$$\begin{cases} u_\alpha u_P = u_H \\ u_\lambda u_P = 1 \\ u_\eta = 1 \\ u_\beta u_H = 1 \\ u_\beta u_H = u_P \\ u_\alpha = 1 \\ u_\alpha u_\gamma u_P = u_H. \end{cases}$$

Solving the equations we find that the only solution is the trivial one, so the model is locally structurally identifiable. It is worth noting that the method based on differential algebra requires 5 pages of analytical calculations to prove the global identifiability property of this model, and that software packages as DAISY [15], COMBOS [16], STRIKE-GOLDD [17, 18] or SIAN [19] are unable to process this in a reasonable computational time, or provide no answer at all.

### 3 Summary Table

In Table 1 we summarize the main results of our analysis.

**Table 1.** Summary of models analyzed in this work: LSI: Local Structurally Identifiable. * Depending on the number of observed variables, it can become LSI. We cannot test Global Structural identifiability.

| Model | Description | Ref. | LSI? |
|---|---|---|---|
| Model 1 | A simple one-dimensional linear model | This work | No |
| Model 2 | Unidentifiable nonlinear model | [1] | No |
| Model 3 | A local structurally identifiable nonlinear model | [1] | Yes |
| Model 4 | A Lotka-Volterra-like model | This work | No* |
| Model 5 | A batch reactor (Michalis-Menten-like) model | [2] | No |
| Model 6 | Goodwin's model | [3] | No |
| Model 6bis | Goodwin's model (all observed) | [3] | No |
| Model 7 | Circadian clock | [3, 4] | No |
| Model 8 (1 observed) | the HIV model with an eclipse phase | [5, 6] | No |
| Model 8bis (all observed) | the HIV model with an eclipse phase | [5, 6] | No |
| Model 8ter (1 observed) | the linear HIV model without an eclipse phase | [5, 6] | No |
| Model 9 | Glycolysis inspired metabolic pathway | [7] | Yes |
| Model 10 | A high dimensional non-linear model | [3, 4] | Yes |
| Model 11 (all parameters unknown) | NF-$\kappa$B signaling pathway | [3, 8] | No |
| Model 11bis (some parameters known) | NF-$\kappa$B signaling pathway | [3, 8] | Yes |
| Model 12 (1 observed) | Pharmacokinetics model | [9] | Yes |
| Model 12bis (2 observed) | Pharmacokinetics model | [9] | Yes |
| Model 13 | Within-host infectious disease | [10] | Yes |
